# Formulation, Characterization and Pharmacokinetic Evaluation of Amorphous Solid Dispersions of Glibenclamide for Bioavailability Enhancement in Wistar Rats

**DOI:** 10.1101/2024.01.31.578259

**Authors:** Khalid Bashir Mir, Nisar Ahmed Khan, Mehraj D. Shah, Alamgir A. Dar

## Abstract

Oral bioavailability of Glibenclamide (Glb) was appreciably improved by forming amorphous solid dispersion (ASD) with Poloxamer-188 (P-188). Poloxamer-188 substantially enhanced the solubility and thereby dissolution rate of the BCS Class-II drug, Glibenclamide (Glb) and simultaneously exhibited better stabilizing effect of amorphous solid dispersion prepared by solvent evaporation method. The physical state of the dispersed Glibenclamide in the polymeric matrix was characterized by differential scanning calorimetry, X-ray diffraction, scanning electron microscope and Fourier Transform Infrared studies. *In vitro* drug release in buffer (pH 7.2) revealed that amorphous solid dispersion at the Glb-P-188 ratio of 1:6 (SDE_4_) improved the dissolution of Glibenclamide 90% within 3 hrs. Pharmacokinetic study of the solid dispersion formulation (SDE_4_) formulation in wistar rats showed that oral bioavailability of the drug was greatly increased as compared to market tablet formulation, Daonil^®^. Solid dispersion formulation (SDE_4_) resulted in approximately two fold higher AUC_0-24_ hrs. Solid dispersion formulation (SDE_4_) formulation was found stable during the study period of six months.

## 1. Introduction

Glibenclamide [1-(4-(2-(5-chloro-2-methoxybenzamido) ethyl) benzene sulphonyl) -3- cyclohexylurea] is an oral hypoglycemic drug used to control blood glucose levels in type 2 diabetic patients. However, Glb is categorized as BCS Class II, thereby it has low aqueous solubility of 0.01886 mg/ml [1, 2]. Therefore, due to poor water solubility, its oral bioavailability is 40% [3–5]. Despite having good pharmacological activity, low oral bioavailability can be major limiting factor for its stoppage in the clinical use since one decade.

Numerous strategies have been employed till now to increase the oral bioavailability of Glb, such as size reduction, the use of surfactants, salt formation, pH adjustments, prodrugs, complexation with beta-cyclodextrins, self-emulsifying formulations, liposomes, Solid-Lipid Nanoparticles, Nanocrystals. [6, 7]. H[8, 9](Martena, Censi, Hoti, Malaj, & Di Martino, 2013; Shah, Parikh, Chavda, & Sheth, 2014)owever the formulations prepared by above mentioned techniques besides having stability issues does not increase the solubility to extraordinary level. So, we formulated the ASD of Glb with the P-188. P-188 is a third generation block co-polymer with good stabilising effect and simultaneously having better solubilizing capacity for a hydrophobic drug candidate like Glb.

A substitute to these strategies is the use of solid dispersions, which can be described as “a group of solids consisting of at least two different components, usually a hydrophilic matrix and a hydrophobic drug, where the drug can be dispersed molecularly or in the amorphous state.”[10]. Solid dispersion is a viable and cost-effective technique to augment bioavailability of poorly water-soluble drugs and also overcomes the limitations of the earlier approaches [6]. The improvement of the dissolution of drugs from solid dispersions is based mainly on three different mechanisms: the wettability of the drug (which is improved by direct contact with the hydrophilic matrix), the reduction in the particle size and increased surface area, and the conversion of the crystalline state to the more soluble amorphous state [11]. Synthetic polymers such as poloxamers, polyvinylpyrrolidone, polyethylene glycols can be used to obtain solid dispersions [12]. Poloxamers has been extensively studied as an efficient carrier in solid dispersions of various drugs and as a result of its ability to retard and inhibit recrystallization of drugs due to its high glass transition temperature. As per available literature, this is the first study reporting the manufacturing of solid dispersions containing Poloxamer-188 and Glibenclamide, which can be an interesting approach to enhance the dissolution profile of a lipophilic drug, employing inexpensive, straightforward and industrially viable technique. Based on this, solid dispersions employing solvent methods and physical mixtures, were prepared, characterized using differential scanning calorimetric, X-ray diffraction, infrared spectroscopy, scanning electron microscopy, solubility and dissolution profiles and evaluated them for bioavailability enhancement by pharmacokinetics studies.

## 2. Materials and Methods

### 2.1 Materials

Glibenclamide was purchased from Sigma-Aldrich. Poloxamer-188, Poloxamer-407, were kindly gifted by Cadila Healthcare Ltd. (Ahmedabad, India). Polyethylene Glycol-4000, Polyethylene Glycol-6000, Polyvinyl Pyrrolidone-K30, Polyvinyl Pyrrolidone-K90, Mannitol, Urea were purchased from Sisco Research Limited (Chandigarh, India). All other chemicals purchased (acetonitrile, glacial acetic acid and methanol) were of HPLC grade. Male wistar rats were purchased from Sher-e-Kashmir University of Agricultural Sciences and Technology (Srinagar, India).

### 2.2 Screening of Carriers

The aqueous solubility of Glibenclamide was tested in different ratios of the aqueous solutions of various carriers (P-188, P-407, PEG-4000, PEG-6000, PVP-K30, PVP-K90, Mannitol and Urea) to select the ideal carrier in an optimum drug-carrier ratio. Phase solubility studies were conducted using the method reported by Higuchi and Connors [13]. An excess quantity of glibenclamide (20 mg) was added to the aqueous solutions of various carriers in different concentrations ranging from 0–6% w/v. The vials were closed and shaken for 48 hours in the orbital-incubator shaker at 100 rpm and 37±0.5°C. The samples were then centrifuged at 3000 rpm for 20 minutes and filtered through a 0.45μm membrane. The filtrates were suitably diluted and analyzed by UV spectrophotometer for the dissolved drug at λ_max_ 291 nm.

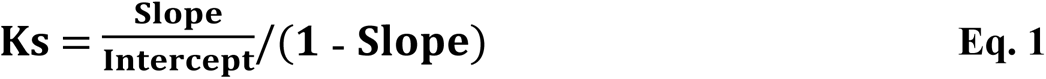

Where, K_s_ is the Apparent Stability Constant

The values of Gibbs free energy of transfer (ΔG°_tr_) of Glibenclamide were computed using Equation:

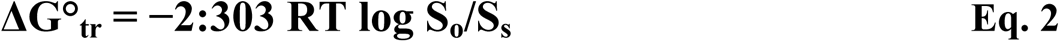

Where, **S_o_** and **S_s_** are the molar solubility of Glibenclamide in aqueous solutions of each carrier to that in pure distilled water, respectively. **T** is absolute temperature and **R** is the universal gas constant.

### 2.3 Preparation of solid dispersions and physical mixtures

#### 2.3.1 Solid dispersions by Solvent Evaporation Method

In this method, the drug and carrier were dissolved in ethanol with constant stirring till a solution is formed and then transferred to a petridish for complete evaporation of solvent at room temperature. The mass was then dried at 25°C temperature in hot air oven, then pulverized and sieved through mesh No. 60. The powdered mass collected was stored in glass vials and kept in a desiccator till further analysis [14].

#### 2.3.2 Physical mixtures by Physical Mixing

In this method, the drug and carrier were mixed using pestle & mortar for 30 minutes and the physical mixtures were sieved through mesh No. 60. The powder mass collected was stored in glass vials and kept in a desiccator till further analysis [15].

### 2.4. Differential Scanning Calorimetry (DSC)

Thermal analysis was performed using differential scanning calorimeter. Under nitrogen flow of 20 ml/min, approximately 8 mg of plain Glibenclamide, plain Poloxamer-188 and selected solid dispersion formulation ‘SDE_4_’ were separately placed in sealed aluminum pans and heated at a scanning rate of 10°C/min from 30°C to 700°C. An empty aluminum pan was used as a reference [16, 17].

### 2.5. Powder X-Ray Diffractometry (PXRD)

Powder X-ray diffraction (XRD) patterns for plain Glibenclamide, plain Poloxamer-188 and selected solid dispersion formulation (SDE_4_) were recorded in diffractometer using target CuK*a* monochromatized radiation, voltage 40 KV, and current 40 mA at ambient temperature. The data was collected in the continuous scan mode using a step size of 0.01° at 2θ/s. The scanned range was 5-90° [18–21].

### 2.6. Scanning Electron Microscopy (SEM)

Morphology studies of plain Glibenclamide, plain Poloxamer-188 and selected solid dispersion formulation ‘SDE_4_’ were conducted in scanning electron microscope. The samples were fixed separately on aluminum stubs using mutual conductive adhesive tape and sputter-coated with a gold layer at 20 mA for 30 s in an ion sputter coater at a pressure of 8–10 Pa prior to the observation at an accelerating voltage of 20 kV [22–24].

### 2.7. Fourier Transform Infrared Spectroscopy (FT-IR)

The interactions studies between drug and carrier in the solid dispersion formulation ‘SDE_4_’ were confirmed by IR spectroscopy, in which the dried samples of plain Glibenclamide, plain Poloxamer-188 and selected solid dispersion formulation (SDE_4_) were separately mixed with potassium bromide and then scanned in the fourier transform infrared spectrophotometer (FT-IR) between 4000 cm^-1^ to 500 cm^-1^[25–29].

### 2.8. Solubility Studies

In this method, an excess amount of plain Glibenclamide (20 mg), physical mixtures and solid dispersions (containing an equivalent amount of Glibenclamide, 20 mg) were taken in 25 ml conical flasks containing 10 ml of phosphate buffer pH 7.2. The flasks were shaken for 24 hours at 100 rpm and temperature of 37±0.5°C in orbital-incubator shaker. The solutions were filtered using 0.45 µm membrane filter. The filtrate obtained was suitably diluted with respective phosphate buffer (pH 7.2) and the amount of drug was estimated using UV spectrophotometer at their relevant λ_max_ 300 nm [30].

### 2.9. Dissolution Studies

The *in-vitro* drug release studies were conducted in USP Type II Dissolution Apparatus using 900 ml of phosphate buffer pH 7.2 at 37±0.5°C with stirring rate of 100 rpm. The plain Glibenclamide (20mg), physical mixtures and solid dispersions in the powdered form (containing an equivalent amount of drug Glibenclamide, 20 mg) were introduced into separate dissolution flasks and the samples of 5 ml volume were withdrawn at predetermined time intervals and filtered through 0.45 µm membrane filter and simultaneously, the dissolution media were replenshised with fresh 5 ml of solvent/buffer to maintain the artificial sink conditions. The filtrate obtained was suitably diluted with the respective solvent/buffer and the amount of drug was estimated using UV spectrophotometer at the relevant λ_max_ 300 nm [31–33].

### 2.10 Comparative dissolution studies

The comparative in-vitro dissolution studies were performed on marketed tablet formulation (Daonil, Sanofi Aventis) and selected solid dispersion formulation ‘SDE_4_’ in phosphate buffer, pH 7.2. The in-vitro drug release studies were conducted in USP type II dissolution apparatus using 900 ml of phosphate buffer, pH 7.2 at 37±0.5°C with stirring rate of 100 rpm. Both the marketed formulation and selected solid dispersion in the powdered form containing an equivalent amount of drug (20 mg) were introduced into separate dissolution flasks and the samples of 5 ml volume were withdrawn at predetermined time intervals and filtered through 0.45 µm membrane filter and simultaneously, the dissolution media were replenshised with fresh 5 ml of phosphate buffer, pH 7.2 to maintain the artificial sink conditions. The filtrates obtained were suitably diluted with phosphate buffer (pH 7.2) and the amount of drug was estimated using UV spectrophotometer at the λ_max_ 300 nm [34–36].

### 2.11 Pharmacokinetic Studies

The animal experiment was approved by Institutional Animal Ethics Committee (IAEC) at the University of Kashmir, Hazratbal, Srinagar, J&K, India under No: F(IAEC-Approval) KU/2017/08 Dated: 16-12-2017. The analytical study was performed at the Research Centre for Residue and Quality Analysis-Sheri Kashmir University of Agricultural Sciences & Technology of Kashmir, Shalimar, Srinagar, J&K, India. Male wistar rats having body weights of approximately 250-300 grams were used. The rats were maintained in a well ventilated animal house with alternating 12-h light/dark cycles at a room temperature of 22±2°C and a relative humidity of 50±10%. The rats were given commercial rat chow and water *ad libitum*. Twenty Four rats were randomly divided into four groups having six rats in each group and the animals were kept fasted for 12 hours prior to start of experiment, but had free access to glucose-water solution throughout the experimental period to prevent any hypoglycemic condition arising due to Glibenclamide dosing. Each group of rats were given a single dose of Glibenclamide equivalent to 10 mg/kg body weight by intra-gastric intubation, i.e., first group received plain Glibenclamide, second group received marketed formulation, (Daonil–Sanofi Aventis) in powdered form, third group received physical mixture, (PM_4_) and fourth group received selected solid dispersion formulation, (SDE_4_) [37–39]. 500µl of blood collected from the orbital venous plexus at the predetermined time intervals (0.5, 1, 1.5, 2, 3, 4, 8, 12, 24 hours) were placed into pre-treated tubes with heparin. The samples were immediately centrifuged at 10,000 rpm for 15 minutes at 4°C and the plasma obtained was refrigerated at -20°C till further analysis. For the assay of glibenclamide, thawed plasma samples (500 µl) were mixed with 25 µl internal standard spiking solution (Flufenamic acid, 40 µg/ml) and vortex mixed with 1000 µl of ethyl acetate followed by centrifugation at 9000 rpm for 15 minutes at 4°C temperature. After centrifugation, the supernatant organic layer was transferred to another glass tube and evaporated to dryness by nitrogen drying at 40 °C under gas pressure of 3 litres/min. The dried residues was reconstituted with 250 µl of the mobile phase and transferred to the HPLC auto-sampler vials.The HPLC system (Agilent Infinity 1200 series) equipped with C_18_ column (250×4.6 mm Agilent 5 μm, Zorbax, SB) and photodiode array detector was used for sample analysis. The Glibenclamide peak was eluted at 9.01 min and that of internal standard (Flufenamic acid) was eluted at 14.88 min with the mobile phase (500 ml Acetonitrile + 500 ml Water pH 4.0 maintained with Glacial Acetic Acid) running at 1 ml/min. The detection wavelength was 230 nm with an injection volume of 20μL.

#### Analysis of Data

The plasma drug concentration–time data for Glibenclamide in rats were analyzed by software (WinNolin, Version 5.2.1) employing a non-compartmental model. The area under the plasma concentration–time curve from zero to the last measurable plasma concentration point (AUC_0–t_) and from zero to infinity (AUC_0-∞_) was calculated by the linear-log trapezoidal method. ANOVA ‘one way’ was used for the statistical comparison of pharmacokinetic parameters between oral dosing of plain Glibenclamide, powdered market formulation (MF), physical mixtures formulation (PM_4_) and selected solid dispersion formulation (SDE_4_).

### 2.12 Percent Practical Yield

Percent practical yield was calculated to find out the efficiency of the method. Solid dispersions were collected and simply weighed to determine practical yield (PY) from the equation (Eq. 3). The study was performed in triplicate [40].

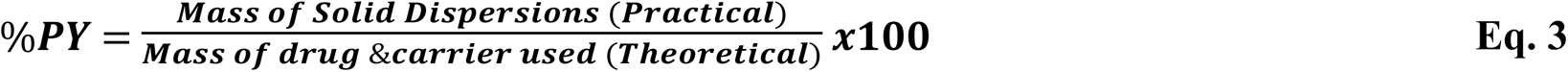

### 2.13. Percent Drug Content

Solid dispersions (containing an equivalent amount of Glibenclamide,20mg) were weighed accurately and dissolved in 10 ml of methanol. The solution was filtered, diluted suitably with methanol and drug content was estimated UV spectrophotometrically at λ_max_ 283 nm. The actual drug content (DC) was calculated using the equation **(Eq. 4)**. The study was performed in triplicate [41].

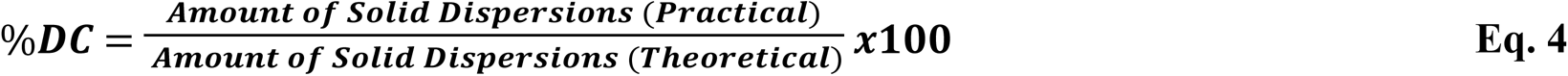

### 2.14. Wetting Time Studies

300 mg of plain Glibenclamide, selected physical mixture (PM_4_), selected solid dispersion formulation (SDE_4_) and marketed tablet formulation (Daonil-Sanofi Aventis) in powdered forms were separately placed in a sintered glass funnel having 33mm internal diameter. The funnel was plunged into beaker containing distilled water, such that the surface of water in beaker was at the same level as the powder in the funnel. 10 mg of methylene blue was layered uniformly on the surface of the powders in the funnel. The time required for wetting of the samples was measured as the time taken to change in color by methylene blue. The study was performed in triplicate.

### 2.15 Stability Studies

Stability studies were conducted to observe the stability of the amorphous form of Glibenclamide in solid dispersion formulation ‘SDE_4_’ during accelerated storage conditions. For accelerated storage conditions, the samples were stored in an environment of 40°C±2°C and 75%RH±5%RH for 6 months (according to the ICH Guidelines-2003 for stability testing of new drug substances and products, Q1AR2) in environmental stability chamber. The samples were analyzed by in-vitro dissolution, drug content and XRD methods at the appropriate time points (0, 1, 3, 6 months). The results obtained were compared with the characterization of the samples at 0 month.

### 2.16 Statistical analysis

All data was expressed as ‘Mean±SD’ (standard deviation). The significant differences were determined statistically using ANOVA ‘one-way’ followed by Tukey-Kramer or Tukey’s Honestly Significant Difference (HSD) Post Hoc Test (multiple comparison test) for identification of individual differences between treatment groups with P<0.05 denoting significance using software (GraphPad Prism, Version 6.0).

## 3. Results and discussion

### 3.1 Screening of Carriers

From the screening of chosen eight carriers (Poloxamer-188, Poloxamer-407, PEG-4000, PEG-6000, PVP-K30, PVP-K90, Mannitol, Urea) by phase solubility studies, Poloxamer-188 showed better potential towards solubility enhancement of Glibenclamide with increase in carrier concentrations from 0-6% (w/v), hence was selected for the preparation of physical mixtures and solid dispersions of Glibenclamide. It is quite clear from the phase solubility profiles (Figure 1), that the highest solubilization of Glibenclamide is achieved by Poloxamer-188.

**Figure 1:**
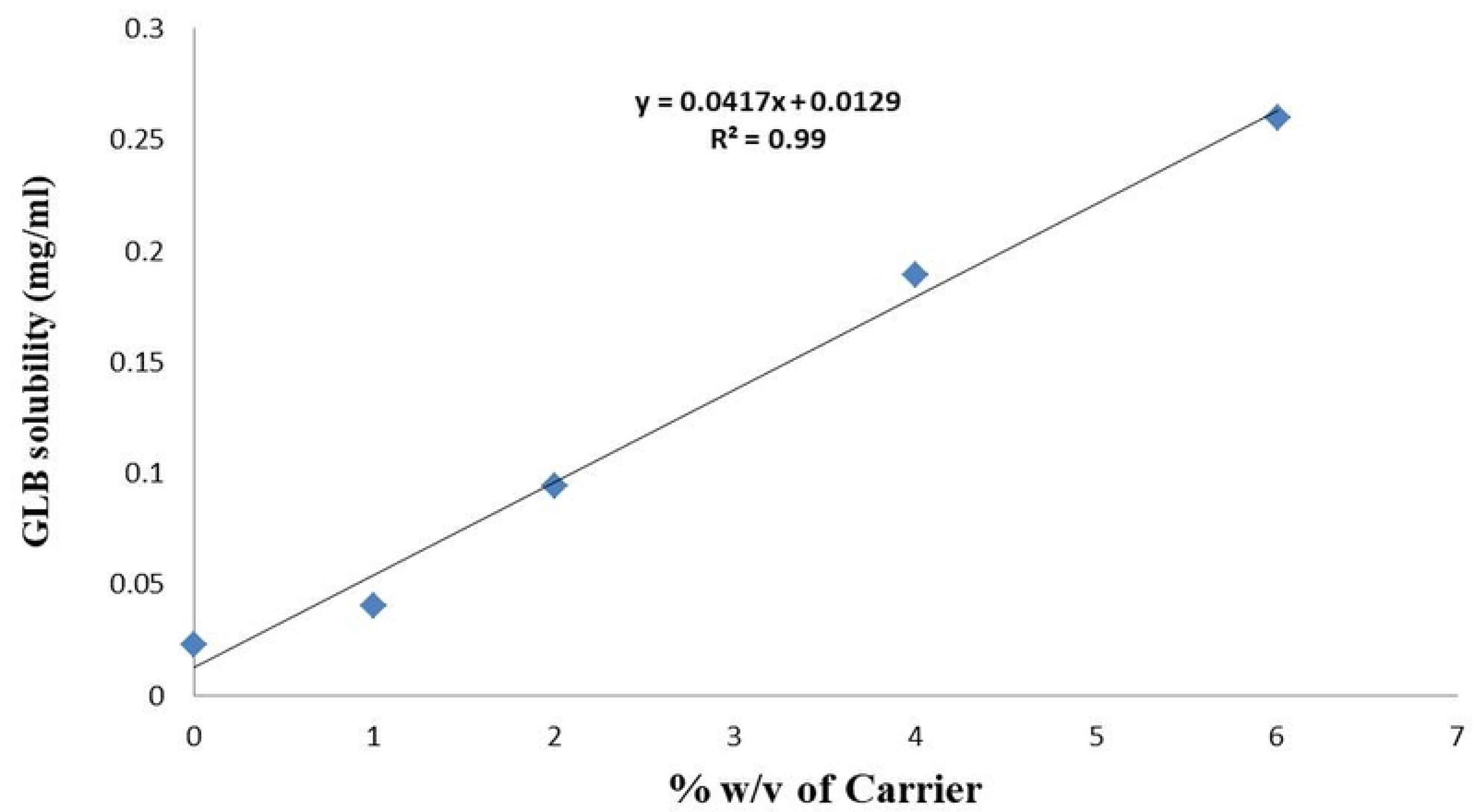
Phase solubility curve for GLB in 0-6% carrier solutions of Poloxamer-188.

Further, the highest potential towards the solubility enhancement of GLB could be attributed to the presence of hydrophilic and hydrophobic components in the P-188, which is mainly responsible for its surface activity/self-emulsifying properties and also, due to favorable critical micelle concentration (CMC) value of poloxamer-188. The Poloxamer-188 has more solubilization and micellization capacity for Glibenclamide than other carriers used.

### 3.2 Thermal Studies by DSC

The thermogram of plain Glibenclamide (Figure 2A) shows a sharp endothermic peak at

**Figure 2:**
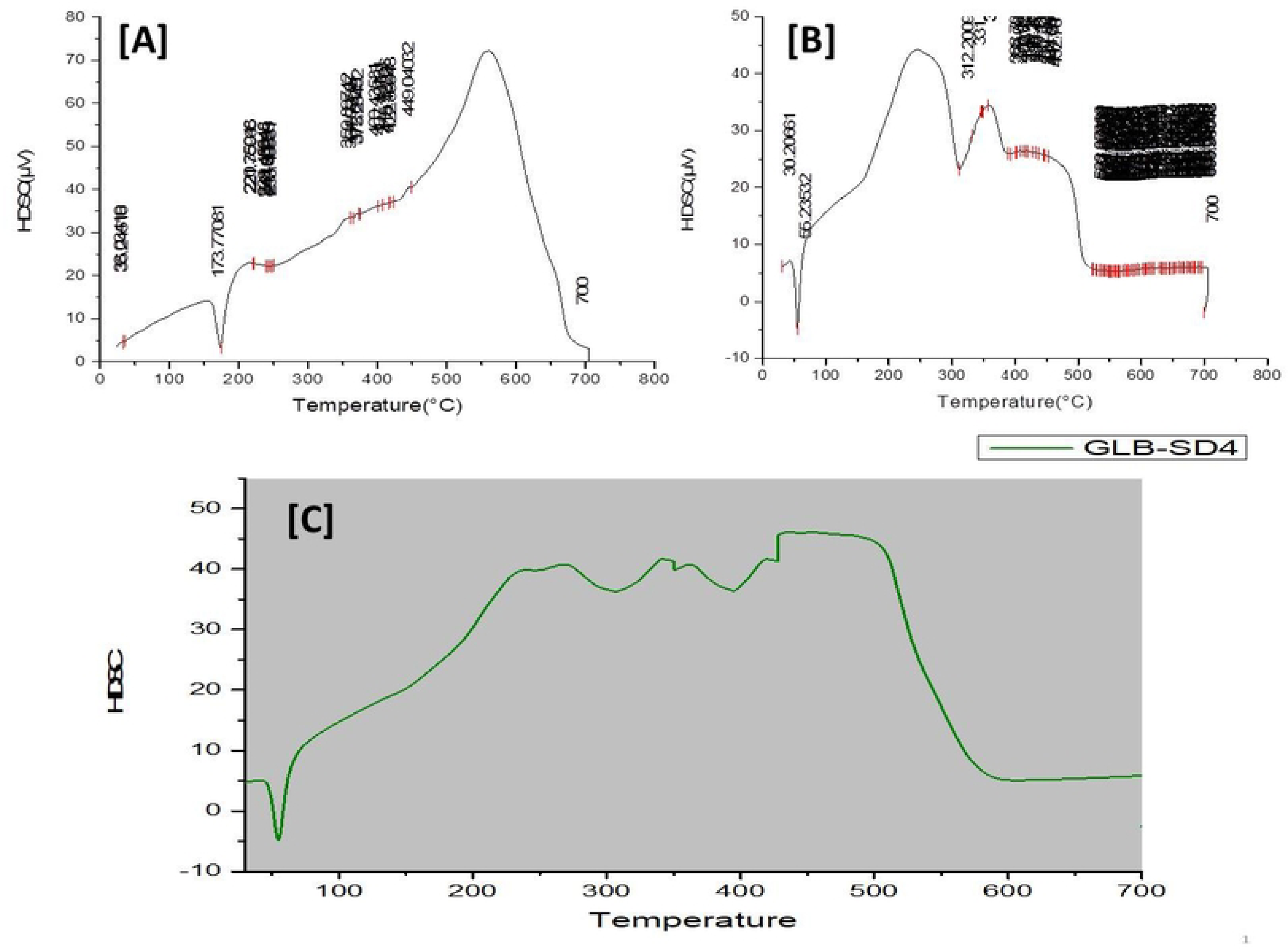
**[A]** DSC thermogram of plain Glibenclamide; **[B]** DSC thermogram of plain Poloxamer-188**; [C]** DSC thermogram of selected solid dispersion ‘SDE_4_’formulation.

173.77°C relative to its melting point, indicating its crystalline nature and its melting point, while the thermogram of Poloxamer-188 (Figure 2B) showed an endothermic peak relative to its melting point at 55.235°C. In the thermogram of the solid dispersion **‘SDE_4_’** formulation (Figure 2C), the melting point of Glibenclamide was not detectable, which could be due to presence of Glibenclamide in amorphous form.

### 3.3 Solid State Characterization by PXRD

The diffractogram of plain Glibenclamide (Figure 3A) showed intensive diffraction peaks, with the most prominent peaks being 11.73°, 16.23°, 18.97°, 19.46°, 21.02°, 22.89°, 23.21, 27.76°, 30.33° and 31.81° suggesting crystalline nature of Glibenclamide, while Poloxamer-188 (Figure 3B ) had an XRD pattern with two prominent diffraction peaks at 19.1° and 23.2°, suggesting the presence of crystalline domains within the carrier. Almost all characteristic peaks of Glibenclamide disappeared in the diffractogram of solid dispersion **‘SDE_4_’**formulationindicating the conversion of the crystalline Glibenclamide to amorphous form. The diffraction spectrum of SDE_4_ (Figure 3C) showed peaks at 19.2° and 23.46°, assigned to the carrier which underwent no alteration in its crystalline nature.

**Figure 3:**
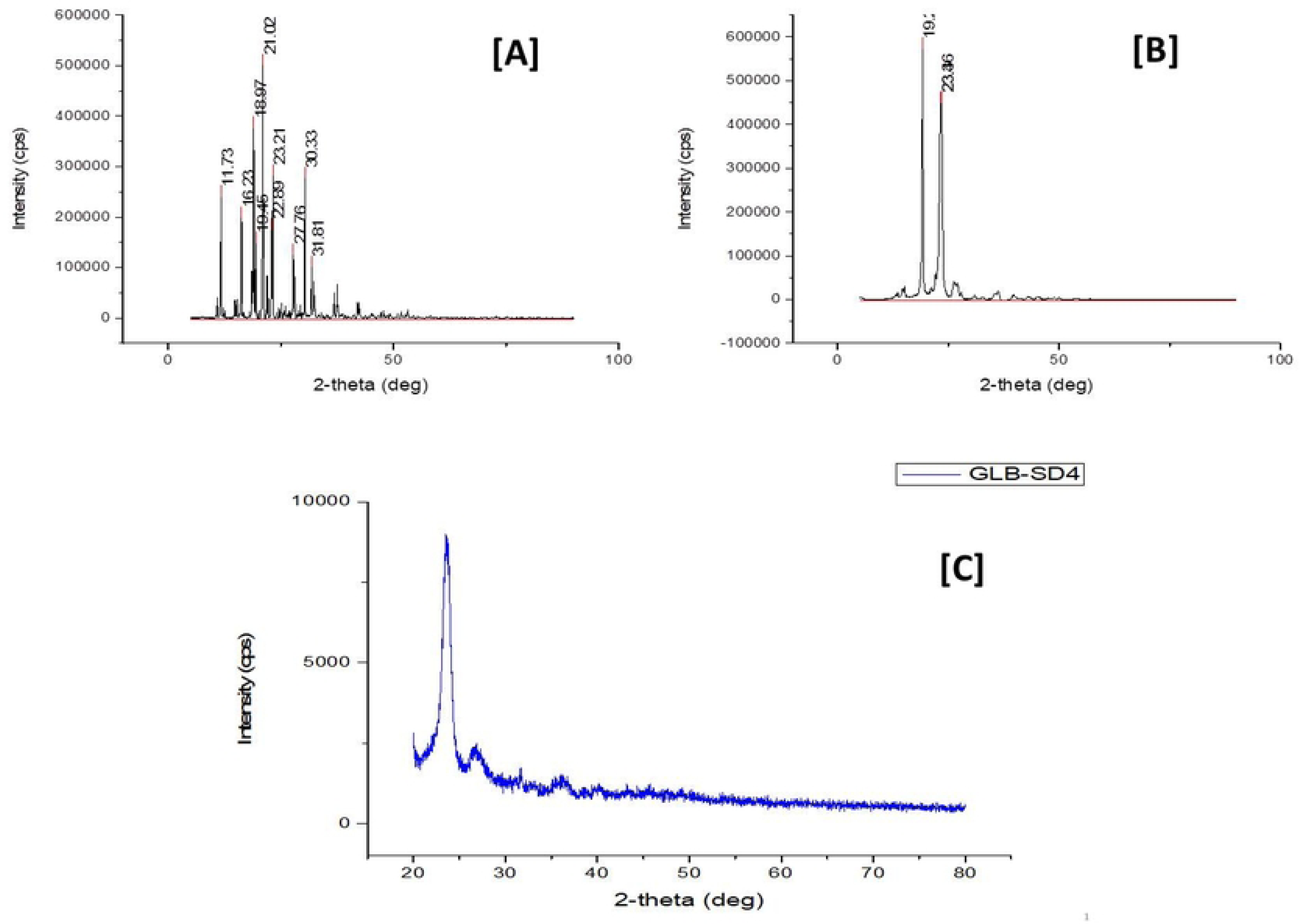
[A] XRD diffractogram of plain Glibenclamide; **[B]** XRD diffractogram of plain Poloxamer-188; [C] XRD diffractogram of selected solid dispersion ‘SDE_4_’formulation.

### 3.4 Surface Morphology Studies by SEM

The plain Glibenclamide appeared to be irregular crystalline solid (Figure 4A), whilePoloxamer-188 was presented as spherical smooth-surfaced particles (Figure 4B). The solid dispersion **‘SDE_4_’**formulation showed wrinkled and irregular surfaced pattern having and the absence of visible Glibenclamide crystals demonstrating the presence of either dissolved Glibenclamide in the carrier or partial conversion of crystalline Glibenclamide into an amorphous form (Figure 4C). The change in surface morphology of Glibenclamide& Poloxamer-188 could be due to process of solubilization and evaporation which can be taken as an indicator of successful formation of solid dispersions. Further, the irregular surface pattern of SDE_4_ could be responsible for enhancement of solubility due to increased surface area.

**Figure 4:**
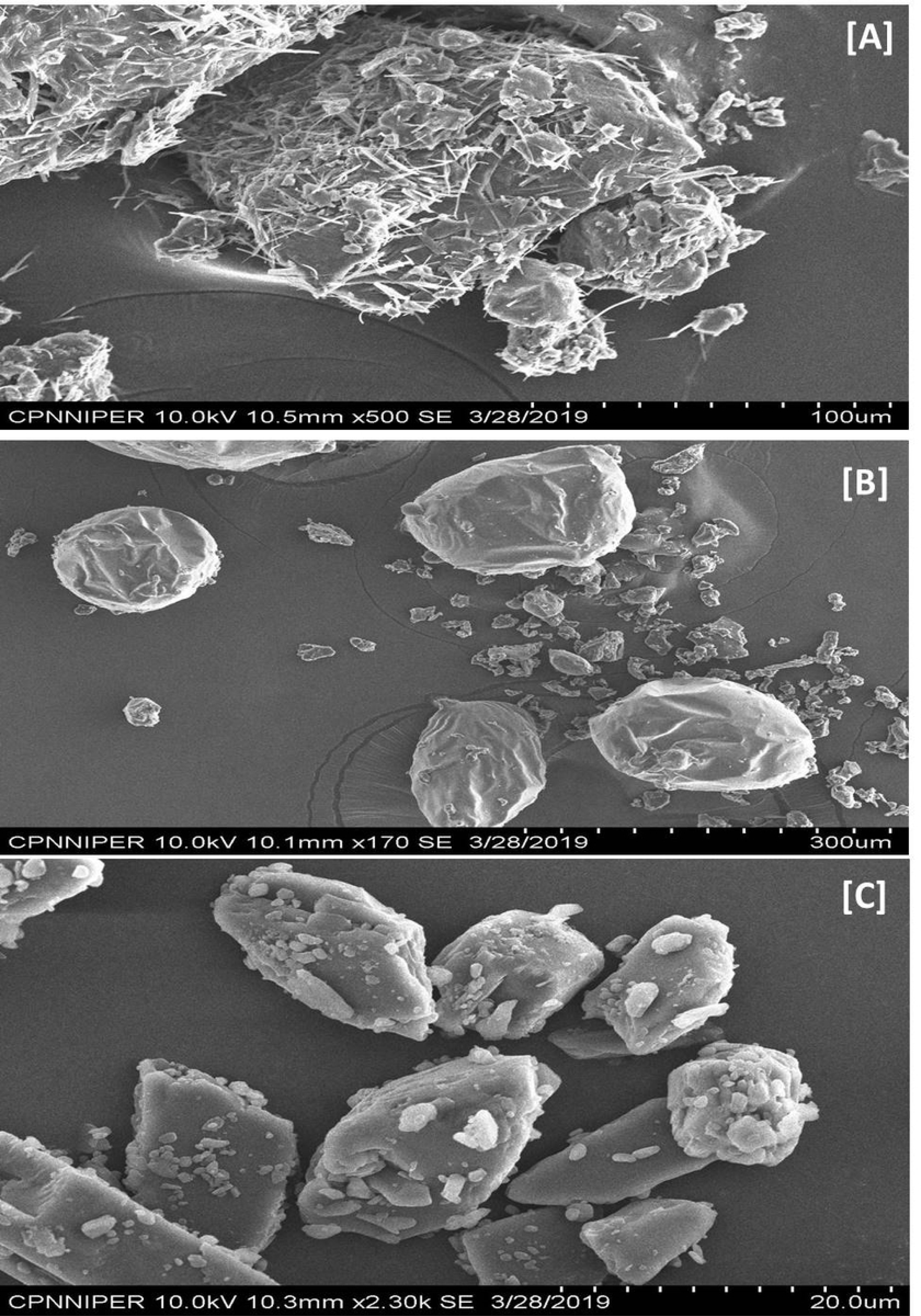
**[A]** XRD diffractogram of selected solid dispersion ‘SDE_4_’formulation; **[B]** SEM microphotograph of plain Poloxamer-188; **[C]** SEM microphotograph of selected solid dispersion ‘SDE_4_’formulation.

### 3.5 Interaction Studies by FT-IR

The FTIR spectrum of plain Glibenclamide **(Figure 5A)** exhibited characteristic peaks, i.e., Amide peaks 3362.1, 3309.9, 3112.3, 2929.7, 2855.1 cm^−1^, Urea N-H stretching peaks 1714.6, 1613.9, 1520.8, 1449.9, 1341.8 cm^−1^ and SO2 stretching vibration peaks 1274.7, 1244.9, 1155.5, 1121.9, 1092.1, 1013.8, 905.7, 741.7, 685.8 cm^−1^. In the IR spectrum of plain Poloxamer-188 **(Figure 5B)**, the characteristic peak at 2877.5, cm^−1^ was observed due to O-H stretching and 1468.6, 1341.8, 1278.5, 1241.2, 1148.0, 1095.8 cm^−1^ were due to C-O stretching and 961.7, 842.4 cm^−1^ were due to bending vibrations respectively. While, the IR spectrum of the SDE_4_ **(Figure 5C)** is very similar to the spectra of the individual components (Glibenclamide & Poloxmer-188), although the intensities were attenuated, probably due to the lower proportion of Glibenclamide in SDE_4_. Also, the absorption peak of N-H of the Glibenclamide got shifted, which could be attributed to the formation of intermolecular hydrogen bonding between Glibenclamide and two terminal OH groups of Poloxamer-188, which may disrupt the crystalline structure of Glibenclamide. Further the lack of any additional peaks in the spectra of ‘SDE_4_’ formulation confirms the absence of any chemical interactions between drug and carrier in the said solid dispersion formulation

**Figure 5:**
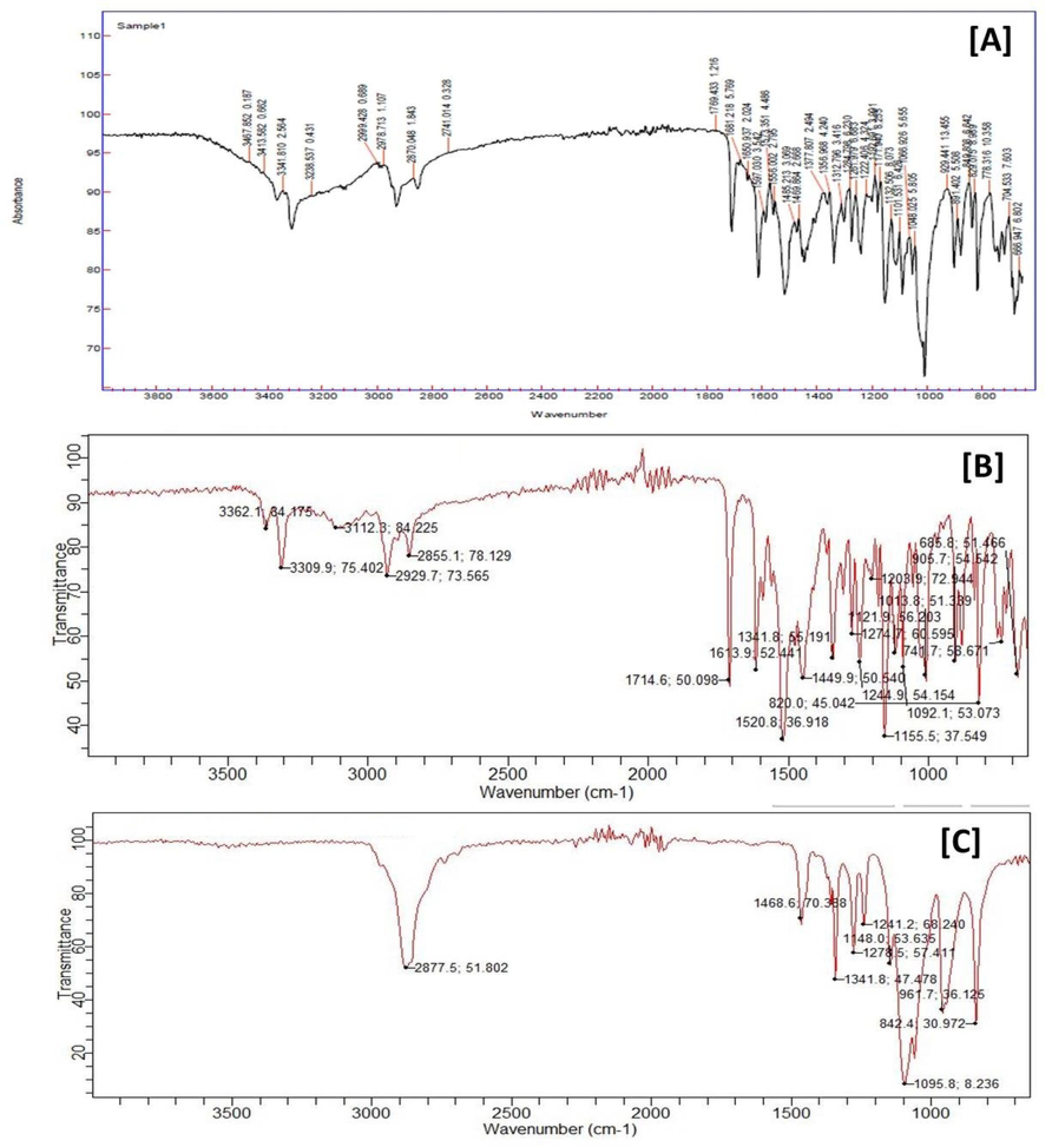
**[A]** FT-IR spectrum of solid dispersion ‘SDE_4_’formulation; **[B]** FT-IR spectrum of plain Glibenclamide; **[C]** FT-IR spectrum of Poloxamer-188.

### 3.5 Solubility Studies

The saturation solubility studies of prepared physical mixtures and solid dispersions were conducted in phosphate buffer, pH 7.2. Out of the 8 formulations, the physical mixtures ‘PM_4_’ and solid dispersions ‘SDE_4_’ showed better solubility enhancement of Glibenclamide in phosphate buffer, pH 7.2, while solid dispersion formulation ‘SDE_4_’ was found to have higher solubility increasing potential in comparison to physical mixture formulation ‘PM_4_’. The solubility enhancement of GLB was found highest by solid dispersion formulation ‘SDE_4_’ than other SD formulations and PM formulations and this could be attributed to the use optimum drug-carrier ratio (1:6 w/w) which has led to improved wettability, higher porosity, reduced size of Glibenclamide (Figure S1).

### 3.6 Dissolution Studies

The dissolution profile of physical mixture ‘PM_4_’ and solid dispersion ‘SDE_4_’ selected on the basis of highest saturation solubility profiles in phosphate buffer, pH 7.2 followed the same trend as observed in the saturation solubility studies (table 1). And, the results indicated that solid dispersion formulation ‘SDE_4_’ showed the highest dissolution rate in phosphate buffer, pH 7.2, and confirmed the highest saturation solubility profile of ‘SDE_4_’ formulation as valid, thereby ‘SDE_4_’ was finalized as the selected solid dispersion formulation for further analysis. This behaviour of SDE_4_ is attributed to the presence of crystalline GLB in the highly energetic amorphous form and also, the complete stabilization of the amorphous GLB by P-188, maintaining the supersaturation over the time by impeding the recrystallization of GLB (Figure S2).

**Table 1:** ***In-vitro* drug release of PM4 & SDE4 in Phosphate Buffer, pH 7.2**

|  | **Cumulative % drug released** |  |
| --- | --- | --- |
| **Time**<br>**(mins)** | **PM<sub>4</sub>** | **SDE<sub>4</sub>** |
| **05** | 9.341 | 29.245 |
| **10** | 17.979 | 41.994 |
| **15** | 33.463 | 58.461 |
| **30** | 37.896 | 71.897 |
| **60** | 40.244 | 82.243 |
| **120** | 43.343 | 93.348 |

### 3.7 Comparative Dissolution Studies

The comparative in-vitro dissolution studies between the selected solid dispersion **‘SDE_4_’** formulation and the marketed formulation **(Daonil-Sanofi Aventis)** depicted that the cumulative % drug release of solid dispersion formulation ‘**SDE_4_’** was found to be 98% which is much higher than the 52% of marketed tablet formulation. As the **‘SDE_4_’** formulation contains the Poloxamer-188, which is a hydrophilic block copolymer carrier with additional surfactant activity and is prepared by solvent evaporation method. So there is better wettability, reduced particle size, higher porosity and the Glibenclamide existed in the amorphous form where no energy is required to break up the crystal lattice during the dissolution process (Figure S3).

### 3.8 Pharmacokinetic Studies

From the pharmacokinetic parameters (Table 2), it was found that the AUC_0–24h_ of the solid dispersion formulation **‘SDE_4_’** (5636 ng.h/ml) represents greater improvement than that of marketed formulation (3747 ng.h/ml) and AUC_0–24h_ of solid dispersion formulation **‘SDE_4_’** was almost 2-fold compared with marketed. It was also found that C_max_ of solid dispersion formulation was 610.0 ng/ml and thus the difference was highly significant (P < 0.05) compared with C_max_ of marketed formulation (287.5ng/ml). The T_max_ of solid dispersion SDE_4_ (1 h) and marketed formulation (3 h) was faster than that of marketed formulation. The MRT values for SDE_4_ (8.69 h) was also significantly different (P < 0.05) from marketed tablet (9.22 h).The relative bioavailability of SDE_4_ with respect to plain Glibenclamide was obtained as 150.41 and 79.1%, respectively. The value of Relative Bioavailability was also significantly different (P < 0.05) from plain Glibenclamide. The hydrophilicity of Glibenclamide increases through solid dispersion technique. All these could explain that the T_1/2_of solid dispersion formulation was shorter than that of marketed formulation. As the plasma concentration and AUC_0–24h_increased with administered solid dispersion **‘SDE_4_’** formulation. The considerable increase in the C_max_ with simultaneous reduction in T_max_inSDE_4_ could be attributed to increased solubility of Glibenclamide in SDE_4_ due to both solubilization effect of Poloxamer-188 on Glibenclamide and conversion of crystalline Glibenclamide to amorphous form. The oral absorption of Glibenclamide from solid dispersion **‘SDE_4_’**formulation resulted in 2-fold enhancement in oral bioavailability in comparison with marketed formulation. The enhancement in oral bioavailability of Glibenclamide from SDE_4_ was possible due to the use of the hydrophilic carrier (Poloxamer-188) in the formulation of SDE_4_ prepared by solvent evaporation method using ethanol as the solvent in comparison with PM_4_, plain Glibenclamide and marketed formulation. Poloxamer-188 is known to enhance the oral bioavailability of lipophilic drugs by enhancing their wettability, surface area, solubility and dissolution rate. Therefore, the significant enhancement in oral bioavailability of Glibenclamide from solid dispersion **‘SDE_4_’**formulation was probably due to the presence of Glibenclamide in amorphous form and Poloxamer-188 also prevents recrystallization of amorphous Glibenclamide back to the initial crystalline state by inhibiting nucleation and crystal growth during in-vivo dissolution. Further, the poloxamer-188 has also a huge role to maintain the supersaturation during in-vivo dissolution.

**Table 2:** Pharmacokinetic parameters.

| **Pharmacokinetic parameters** | **Plain GLB**<br>**(Mean ±SD)**<br>**n=6** | **MF Daonil (Mean**<br>**±SD) n=6** | **PM<sub>4</sub>**<br>**(Mean ±SD)**<br>**n=6** | **SDE<sub>4</sub>**<br>**(Mean ±SD)**<br>**n=6** |
| --- | --- | --- | --- | --- |
| C<sub>max</sub> (ng/ml) | 238.5 ± 0.54 | 287.5 ± 0.41 | 363.5 ± 0.93 | 610.0 ± 0.93 |
| T<sub>max</sub> (h) | 4.07 ± 0.017 | 3.06 ± 0.020 | 2.06 ± 0.013 | 1.07 ± 0.026 |
| AUC<sub>0-24</sub> (ng•hml<sup>-1</sup>) | 2807 ± 23.97 | 3747 ± 22.19 | 3592 ± 11.27 | 5636 ± 19.84 |
| K<sub>el</sub> (h<sup>-1</sup>) | 0.075 ± 0.002 | 0.197 ± 0.005 | 0.437 ± 0.011 | 0.497 ± 0.005 |
| t<sub>1/2</sub> (h) | 9.24 ± 0.006 | 3.517 ± 0.004 | 1.585 ± 0.005 | 1.394 ± 0.003 |
| MRT (h) | 9.23 ± 0.009 | 9.22 ± 0.007 | 9.42 ± 0.007 | 8.69 ± 0.006 |
| Relative<br>Bioavailability (%) | 79.91 | — | 95.86 | 150.41 |

### 3.9 Percent Practical Yield

The percent practical yield of solid dispersions prepared by solvent evaporation method was found to be in the range of 98.12-98.76%. This better yield found with the solvent evaporation method indicates that the method is efficient.

### 3.10 Percent Drug Content

Percent drug content of Glibenclamide in PMs and SDs was found to be in the range of 98.01 to 99.98%. This drug content indicates that the poloxamer-188 has the highest drug loading capacity for Glibenclamide.

### 3.11 Wetting Time Studies

The wetting time of the solid dispersion formulation ‘SDE_4_’ was found to be 15 secs, which is much shorter than those of physical mixture formulation ‘PM_4_’ (45 secs), marketed tablet formulation ‘Daonil’ (47 secs) and plain GLB (50 secs). This short wetting time by SDE_4_ indicates the improvement in wetting capacity of Glibenclamide in solid dispersion formulation due to Polxamer-188 in highest ratio (Fig. S4).

### 3.12 Stability Studies

The results presented in table 3 acquired revealed that no significant differences in the dissolution profile and drug content were observed throughout the 6 month study period. Also, diffractograms from XRD obtained were similar, as those obtained initially, indicating no change in the physical state of the amorphous Glibenclamide in the **‘SDE_4_’** during the 6 month study period (Figure S5-S10).

**Table 3:**
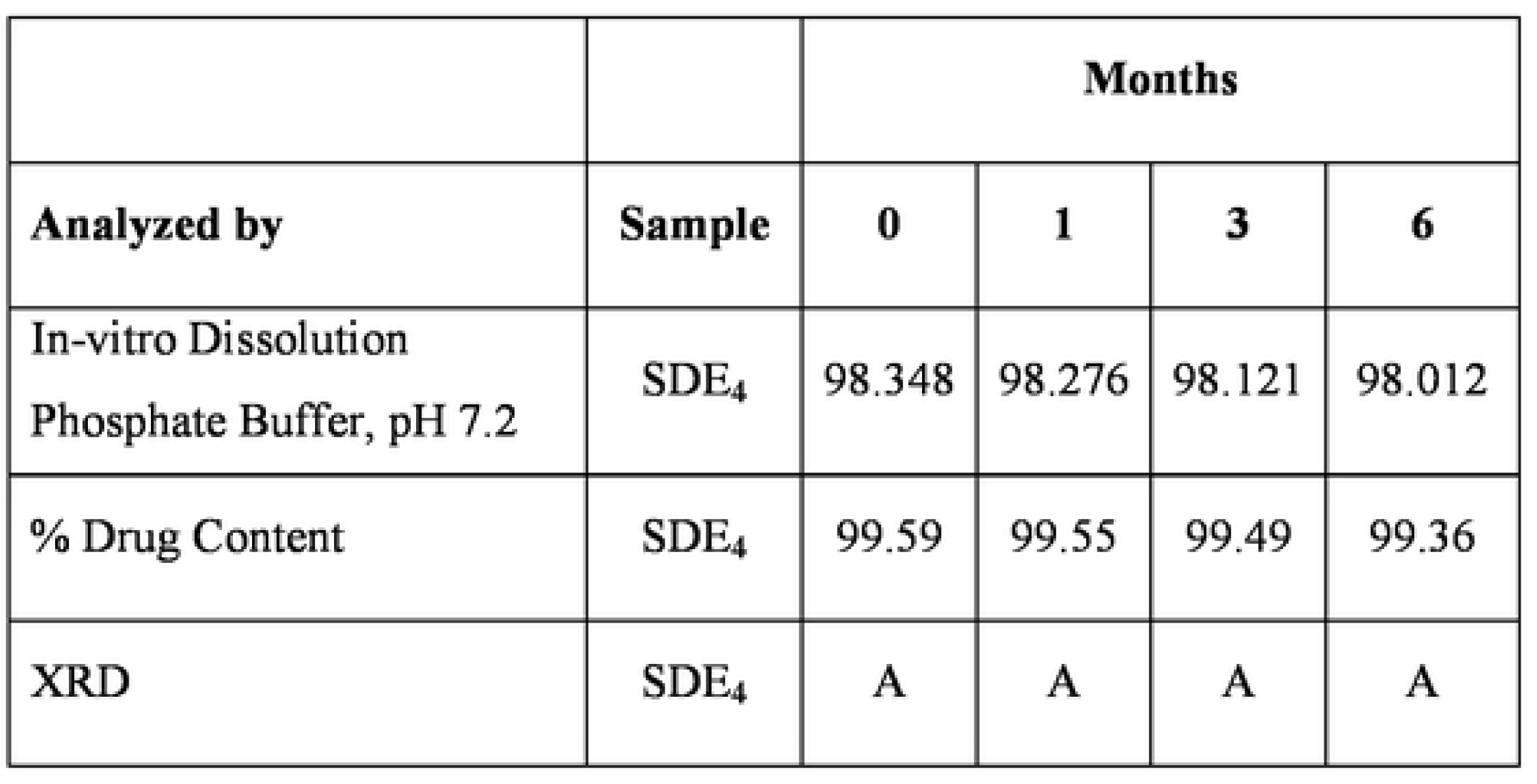
Stability profile under conditions of 40 °C/75% RH.

### 3.16 Statistical Studies

The **‘p’** value obtained from different formulations is <0.05, which showed the differences among the formulations are significant.

## 4. Conclusion

The present research work describes that the solubility, dissolution rate and oral bioavailability of the poorly soluble BCS Class II drug, Glibenclamide was enhanced by the solid dispersion technology. The selected solid dispersion formulation**‘SDE_4_’**of Glibenclamide with Poloxamer-188 (3^rd^ generation carrier) prepared in the 1:6 drug:carrier ratio (w/w) by ‘Solvent Evaporation Method’ showed the highest potential for increased solubility (20 times in Phosphate Buffer, pH 7.2), high in-vitro dissolution rate (3 times in Phosphate Buffer, pH 7.2) and significant increase in oral bioavailability (2-folds in rats) of Glibenclamide than market formulation (Daonil). Besides other factors, like enhanced wetting of the Glibenclamide, size reduction of the Glibenclamide to the molecular level, enhanced particle porosity and less particle agglomeration, the presence of Glibenclamide in high energy amorphous form in the **‘SDE_4_’** formulation was mainly responsible for this enhancement. The optimized solid dispersion formulation**‘SDE_4_’** produced was stable for the study period of 6 months due to absence of any chemical drug-carrier interaction and enormous capability of the Poloxamer-188 to inhibit the nucleation and crystal growth, thus preventing drug recrystallization during both dissolution as well as storage.

## List of Abbreviations

HPLC: High performance liquid chromatography
FT-IR: Fourier Transform Infrared
SEM: scanning electron microscope
Glb: Glibenclamide
SDE_4_: Solid dispersion formulation
ASD: amorphous solid dispersion
P-188: Poloxamer-188
DSC: Differential Scanning Calorimetry

## Data availability statement

Data will be made available on request.

## Additional information

No additional information is available for this paper.

## Declaration of competing interest

The authors declare that they have no known competing financial interests or personal relationships that could have appeared to influence the work reported in this paper.

## Acknowledgements

The authors acknowledge the support of Lark Labs. (New Delhi, India) and Cadila Pharmaceuticals (Ahmedabad, India) for providing gift samples of GLB and Poloxamer-188, respectively. Khalid Bashir Mir is thankful to DST-INSPIRE, New Delhi, India for financial support in the form of Senior Research Fellowship (SRF). The authors are also thankful to Research Centre for Residue and Quality Analysis, Sher-e-Kashmir University of Agricultural Sciences & Technology of Kashmir, Shalimar Campus, Srinagar, J&K, India for providing analytical facility to carry the research study.

## Notes

### Competing Interest Statement

The authors have declared no competing interest.

## References

1. Chauhan B, Shimpi S, Paradkar A. Preparation and evaluation of glibenclamide-polyglycolized glycerides solid dispersions with silicon dioxide by spray drying technique. European journal of pharmaceutical sciences. 2005;26(2):219–30.

2. Elbary AA, Salem H, Maher M. “In vitro and in vivo” Evaluation of Glibenclamide using Surface Solid Dispersion (SSD) Approach. British Journal of Pharmacology and Toxicology. 2011;2(1):51–62.

3. Elbahwy IA, Ibrahim HM, Ismael HR, Kasem AA. Enhancing bioavailability and controlling the release of glibenclamide from optimized solid lipid nanoparticles. Journal of Drug Delivery Science and Technology. 2017;38:78–89.

4. Maiti S, Mukherjee S, Datta R. Core–shell nano-biomaterials for controlled oral delivery and pharmacodynamic activity of glibenclamide. International journal of biological macromolecules. 2014;70:20–5.

5. Lucio D, Irache JM, Font M, Martínez-Ohárriz MC. Nanoaggregation of inclusion complexes of glibenclamide with cyclodextrins. International journal of pharmaceutics. 2017;519(1-2):263–71.

6. Maulvi FA, Dalwadi SJ, Thakkar VT, Soni TG, Gohel MC, Gandhi TR. Improvement of dissolution rate of aceclofenac by solid dispersion technique. Powder technology. 2011;207(1-3):47–54.

7. Gursoy RN, Benita S. Self-emulsifying drug delivery systems (SEDDS) for improved oral delivery of lipophilic drugs. Biomedicine & pharmacotherapy. 2004;58(3):173–82.

8. Shah SR, Parikh RH, Chavda JR, Sheth NR. Glibenclamide nanocrystals for bioavailability enhancement: formulation design, process optimization, and pharmacodynamic evaluation. Journal of Pharmaceutical Innovation. 2014;9(3):227–37.

9. Martena V, Censi R, Hoti E, Malaj L, Di Martino P. Preparation of glibenclamide nanocrystals by a simple laboratory scale ultra cryo-milling. Journal of nanoparticle research. 2013;15(6):1712.

10. Ito A, Watanabe T, Yada S, Hamaura T, Nakagami H, Higashi K, et al. Prediction of recrystallization behavior of troglitazone/polyvinylpyrrolidone solid dispersion by solid-state NMR. International journal of pharmaceutics. 2010;383(1-2):18–23.

11. De Waard H, Hinrichs W, Visser M, Bologna C, Frijlink H. Unexpected differences in dissolution behavior of tablets prepared from solid dispersions with a surfactant physically mixed or incorporated. International journal of pharmaceutics. 2008;349(1-2):66–73.

12. Dangprasirt P, Ritthidej G. Development of diclofenac sodium controlled release solid dispersions by spray drying using optimization strategy I. Powder formulation. Drug development and industrial pharmacy. 1995;21(20):2323–37.

13. Higuchi T. A phase solubility technique. Adv Anal Chem Instrum. 1965;4:117–211.

14. Fouad SA, Malaak FA, El-Nabarawi MA, Abu Zeid K, Ghoneim AM. Preparation of solid dispersion systems for enhanced dissolution of poorly water soluble diacerein: In-vitro evaluation, optimization and physiologically based pharmacokinetic modeling. PLoS One. 2021;16(1):e0245482.

15. Ramadhani N, Shabir M, McConville C. Preparation and characterisation of Kolliphor® P 188 and P 237 solid dispersion oral tablets containing the poorly water soluble drug disulfiram. International journal of pharmaceutics. 2014;475(1-2):514–22.

16. Parmar KR, Shah SR, Sheth NR. Preparation, characterization, and in vitro evaluation of ezetimibe binary solid dispersions with poloxamer 407 and PVP K30. Journal of Pharmaceutical Innovation. 2011;6:107–14.

17. Barmpalexis P, Kachrimanis K, Georgarakis E. Physicochemical characterization of nimodipine– polyethylene glycol solid dispersion systems. Drug Development and Industrial Pharmacy. 2014;40(7):886–95.

18. Kim NA, Choi DH, Lim JY, Kim KH, Lim DG, Lee E, et al. Investigation of polymeric excipients for dutasteride solid dispersion and its physicochemical characterization. Archives of pharmacal research. 2014;37:214–24.

19. Obaidat RM, AlTaani B, Ailabouni A. Effect of different polymeric dispersions on in-vitro dissolution rate and stability of celecoxib class II drug. Journal of Polymer Research. 2017;24:1–14.

20. Dar AA, Jan I, Wani AA, Mubashir S, Sofi KA, Sofi JA, et al. Risk assessment, dissipation behavior and persistence of quinalphos in/on green pea by gas chromatography with electron capture detector. Journal of separation science. 2018;41(11):2380–5.

21. Jan I, Dar AA, Mubashir S, Alam Wani A, Mukhtar M, Sofi KA, et al. Quantification, dissipation behavior and risk assessment of ethion in green pea by gas chromatography-electron capture detector. Journal of separation science. 2018;41(9):1990–4.

22. Ellenberger DJ, Miller DA, Kucera SU, Williams III RO. Improved vemurafenib dissolution and pharmacokinetics as an amorphous solid dispersion produced by KinetiSol® processing. AAPS PharmSciTech. 2018;19(5):1957–70.

23. Teixeira C, Mendonça L, Bergamaschi M, Queiroz RHC, Souza GEPd, Antunes LMG, et al. Microparticles containing curcumin solid dispersion: stability, bioavailability and anti-inflammatory activity. AAPS PharmSciTech. 2016;17:252–61.

24. Guenther P, Guenther M, Ringle CM, Zaefarian G, Cartwright S. Improving PLS-SEM use for business marketing research. Industrial Marketing Management. 2023;111:127–42.

25. Tiernan H, Byrne B, Kazarian SG. ATR-FTIR spectroscopy and spectroscopic imaging for the analysis of biopharmaceuticals. Spectrochimica Acta Part A: Molecular and Biomolecular Spectroscopy. 2020;241:118636.

26. Khan S, Madni A, Rahim MA, Shah H, Jabar A, Khan MM, et al. Enhanced in vitro release and permeability of glibenclamide by proliposomes: Development, characterization and histopathological evaluation. Journal of Drug Delivery Science and Technology. 2021;63:102450.

27. Silva Filho SF, Pereira AC, Sarraguça JM, Sarraguça MC, Lopes J, de Freitas Façanha Filho P, et al. Synthesis of a glibenclamide cocrystal: full spectroscopic and thermal characterization. Journal of Pharmaceutical Sciences. 2018;107(6):1597–604.

28. Ahmad Dar A, Sangwan P, Kumar A. Chromatography: An important tool for drug discovery. Journal of separation science. 2020;43(1):105–19.

29. Dar AA, Dangroo NA, Raina A, Qayum A, Singh S, Kumar A, et al. Biologically active xanthones from Codonopsis ovata. Phytochemistry. 2016;132:102–8.

30. Zhang X, Xing H, Zhao Y, Ma Z. Pharmaceutical dispersion techniques for dissolution and bioavailability enhancement of poorly water-soluble drugs. Pharmaceutics. 2018;10(3):74.

31. Guan J, Jin L, Liu Q, Xu H, Wu H, Zhang X, et al. Exploration of supersaturable lacidipine ternary amorphous solid dispersion for enhanced dissolution and in vivo absorption. European Journal of Pharmaceutical Sciences. 2019;139:105043.

32. Dar AA, Sangwan PL, Singh N, Kumar A. Method validation and simultaneous quantification of five triterpenoids from Codonopsis ovata by high-performance thin-layer chromatography. JPC-Journal of Planar Chromatography-Modern TLC. 2019;32(3):251–6.

33. Singh N, Abrol V, Parihar S, Kumar S, Khanum G, Mir JM, et al. Design, Synthesis, Molecular Docking, and In Vitro Antibacterial Evaluation of Benzotriazole-Based β-Amino Alcohols and Their Corresponding 1, 3-Oxazolidines. ACS omega. 2023;8(44):41960–8.

34. Alzahrani A, Nyavanandi D, Mandati P, Youssef AAA, Narala S, Bandari S, et al. A systematic and robust assessment of hot-melt extrusion-based amorphous solid dispersions: Theoretical prediction to practical implementation. International journal of pharmaceutics. 2022;624:121951.

35. Dar AA, Sangwan PL, Khan I, Gupta N, Qaudri A, Tasduq SA, et al. Simultaneous quantification of eight bioactive secondary metabolites from Codonopsis ovata by validated high performance thin layer chromatography and their antioxidant profile. Journal of pharmaceutical and biomedical analysis. 2014;100:300–8.

36. Dar AA, Rath SK, Qaudri A, Singh B, Tasduq SA, Kumar A, et al. Isolation, cytotoxic evaluation, and simultaneous quantification of eight bioactive secondary metabolites from Cicer microphyllum by high-performance thin-layer chromatography. Journal of separation science. 2015;38(23):4021–8.

37. Mir KB, Abrol V, Wani TU, Jan I, Singh N, Khan NA, et al. Validation and development of RP-HPLC method for quantification of glibenclamide in rat plasma and its application to pharmacokinetic studies in wistar rats. Heliyon. 2023;9(11).

38. Wani TU, Mir KB, Raina A, Dar AA, Jan I, Khan NA, et al. Simultaneous quantification of Losartan Potassium and its Active Metabolite, EXP3174 in Rabbit Plasma by Validated HPLC-PDA. Biomedical Chromatography. 2023:e5645.

39. Dar AA, Abrol V, Singh N, Gashash EA, Dar SA. Recent bioanalytical methods for the isolation of bioactive natural products from genus Codonopsis. Phytochemical Analysis. 2023;34(5):491–506. doi: 10.1002/pca.3253.

40. Pardhi VP, Jain K. Impact of binary/ternary solid dispersion utilizing poloxamer 188 and TPGS to improve pharmaceutical attributes of bedaquiline fumarate. Journal of Drug Delivery Science and Technology. 2021;62:102349.

41. Tran P, Pyo Y-C, Kim D-H, Lee S-E, Kim J-K, Park J-S. Overview of the manufacturing methods of solid dispersion technology for improving the solubility of poorly water-soluble drugs and application to anticancer drugs. Pharmaceutics. 2019;11(3):132.

